# Ketamine effects on anxiety and fear-related behaviors: current literature evidence and new findings

**DOI:** 10.1101/793398

**Authors:** Gabriela P. Silote, Sabrina F.S. de Oliveira, Deidiane E. Ribeiro, Mayara S. Machado, Roberto Andreatini, Sâmia R. L. Joca, Vanessa Beijamini

**Affiliations:** Biochemistry and Pharmacology Graduate Program, Federal University of Espirito Santo, Vitoria, ES, Brazil; Department of Pharmaceutical Sciences, Health Science Center, Federal University of Espirito Santo, Vitoria, ES, Brazil; Pharmaceutical Sciences Graduate Program, Health Sciences Center, Federal University of Espirito Santo, Vitoria, ES, Brazil; Department of Physics and Chemistry, School of Pharmaceutical Sciences, University of São Paulo, Ribeirão Preto, SP, Brazil; Department of Pharmacology, Federal University of Paraná, Curitiba, PR, Brazil; Aarhus Institute of Advanced Studies (AIAS), Aarhus University, Denmark

**Author notes:** Corresponding Author: Vanessa Beijamini Departamento de Ciências Farmacêuticas Centro de Ciências da Saúde, UFES Av. Marechal Campos, 1468, Maruipe Vitoria, ES, 29043-900, Brazil.

**Keywords:** Ketamine, MK-801, anxiety, fear, panic, elevated T-maze, animal models

## Abstract

Ketamine, a non-competitive N-methyl-D-aspartate (NMDA) receptor antagonist, presents rapid and sustained antidepressant effect in clinical and preclinical studies. Regarding ketamine effects on anxiety, there is a widespread discordance among pre-clinical studies. To address this issue, the present study reviewed the literature (electronic database MEDLINE) to summarize the profile of ketamine effects in animal tests of anxiety/fear. We found that ketamine anxiety/fear-related effects may depend on the anxiety paradigm, schedule of ketamine administration and tested species. Moreover, there was no report of ketamine effects in animal tests of fear related to panic disorder (PD). Based on that finding, we evaluated if treatment with ketamine and another NMDA antagonist, MK-801, would induce acute and sustained (24 hours later) anxiolytic and/or panicolytic-like effects in animals exposed to the elevated T-maze (ETM). The ETM evaluates, in the same animal, conflict-evoked and fear behaviors, which are related, respectively, to generalized anxiety disorder and PD. Male Wistar rats were systemically treated with racemic ketamine (10, 30 and 80 mg/kg) or MK-801 (0.05 and 0.1 mg/kg) and tested in the ETM in the same day or 24 hours after their administration. Ketamine did not affect the behavioral tasks performed in the ETM acutely or 24 h later. MK-801 impaired inhibitory avoidance in the ETM only at 45 min post-injection, suggesting a rapid but not sustained anxiolytic-like effect. Altogether our results suggest that ketamine might have mixed effects in anxiety tests while it does not affect panic-related behaviors.

**Highlights:**

- Ketamine induces mixed effects in animal anxiety tests
- Few studies investigated the individual effects of S-ketamine in anxiety/fear tests
- None study evaluated the effects of R-Ketamine on anxiety/fear-related behaviors
- Systemic ketamine does not affect panic-like behaviors in the elevated T-maze

## 1. Introduction

Anxiety disorders are the most common mental disorders affecting the world population (Bandelow and Michaelis, 2015; Craske et al., 2017; Stein et al., 2017). Anxiety is also a frequent symptom in many mental disorders (Millan et al., 2015). Antidepressants and benzodiazepines, drugs that affect the serotonergic and GABAergic neurotransmission, respectively, are commonly used to treat anxiety disorders such as generalized anxiety disorder (GAD) and panic disorder (PD) (Bandelow et al., 2017; Craske et al., 2017; Koen and Stein, 2011; Nash and Nutt, 2005). However, these drugs present several limitations that limit their use (Bandelow et al., 2017; Blier and Abbott, 2001; Nash and Nutt, 2005) and drive the search for other molecular targets and new drugs.

A plenty body of evidence from clinical and preclinical studies suggests the involvement of glutamatergic neurotransmission in anxiety disorders (Averill et al., 2017; Griebel and Holmes, 2013; Harvey and Shahid, 2012; Riaza Bermudo-Soriano et al., 2012). For instance, adolescent patients diagnosed with GAD showed positive correlation between glutamatergic tone and severity of anxiety symptoms (Strawn et al., 2013) whereas patients with social anxiety showed an increase in glutamate/creatine ratio in the anterior cingulate cortex (Phan et al., 2005). In particular, the therapeutic potential of a variety of negative modulators of glutamate transmission has been described over the last years (Ferraguti, 2018; Griebel and Holmes, 2013; Murrough et al., 2015).

Ketamine, a dissociative anesthesic acting as a non-competitive NMDA antagonist receptor, presents promising rapid and transient antidepressant effect in clinical and pre-clinical studies, which can last for up to 14 days (Berman et al., 2000; Diazgranados et al., 2010; Li et al., 2011; Maeng et al., 2008; McCloud et al., 2015; Murrough et al., 2013; Newport et al., 2015). Besides that, patients with major depressive disorder have shown significant reduction of anxiety symptoms after ketamine infusion (Salvadore et al., 2009; Zarate Jr et al., 2006). In contrast, few studies reported the effect of ketamine in patients suffering from anxiety disorders (Banov et al., 2019; Glue et al., 2018, 2017; Irwin et al., 2013; Mohammad Shadli et al., 2018; Taylor et al., 2018). For example, ketamine induced rapid and sustained anxiolytic effect in patients suffering from refractory GAD and/or social anxiety disorder, who were not depressed (Glue et al., 2017). In a case report, ketamine produced a sustained resolution of PD, agoraphobia, and GAD after a single infusion in a patient (Ray and Kious, 2016).

There is a widespread discordance among pre-clinical studies addressing the effects of ketamine on anxiety/fear-related behaviors. Additionally, narrative or systematic reviews on ketamine effects in animal tests of anxiety/fear are lacking. In this context, the current study comprehensively reviewed the literature to summarize the effect profile of ketamine on tests of anxiety/fear-related behaviors to improve our understanding about this topic. We found that ketamine effects on anxiety/fear-related behaviors may depend on several factors, as the type of anxiety evoked by the paradigm, schedule of ketamine administration and tested species/sex. Besides that, we also found that the majority of studies evaluated ketamine effects in animal models related to GAD whereas none study evaluated ketamine effects in animal models related to PD.

Based on the aforementioned considerations, another aim of our study was to investigate if ketamine would induce a rapid and sustained anxiolytic and/or panicolytic-like effect in rats exposed to the elevated T-Maze (ETM). The ETM evaluates, in the same animal, conflict-evoked and innate anxiety behaviors which are related, respectively, to GAD and PD (Moreira et al., 2013; Zangrossi Jr and Graeff, 2014). To this end, we compared the effects of a single injection of ketamine or MK-801 (an NMDA antagonist, also call dizocilpine; Huettner and Bean, 1988; Wong et al., 1986) in the inhibitory avoidance and escape responses generated by the ETM acutely and 24 hours after their administration.

## 2. Material and methods

### 2.1 Literature review

We performed a comprehensive literature review to summarize ketamine effects in studies that evaluate anxiety/fear-related defensive behaviors. We performed a preliminary search in the electronic database MEDLINE and found out that some studies that investigated the antidepressant-like effect of ketamine also evaluated its anxiety-related effects as a secondary outcome. Thus, we decide to include, in our search, terms related to depression. Although the post-traumatic stress disorder (PTSD) had been removed from the anxiety disorder category of the Diagnostic and Statistical Manual of Mental Disorders, Fifth Edition (DSM-5), patients’ suffering from PTSD frequently exhibit signs of anxiety. Consequently, we also included search terms related to PTSD.

Thus, the studies were searched in the electronic database MEDLINE via the search engine PubMed® using the following Medical Subject Headings (MeSH®) terms as descriptors: ((((“Ketamine”[Mesh]) AND (((((((((((((“Depression”[Mesh]) OR “Depressive Disorder”[Mesh:NoExp]) OR “Depressive Disorder, Major”[Mesh]) OR “Depressive Disorder, Treatment-Resistant”[Mesh]) OR “Fear”[Mesh:NoExp]) OR “Panic”[Mesh]) OR “Anxiety”[Mesh:NoExp]) OR “Performance Anxiety”[Mesh]) OR “Stress Disorders, Traumatic”[Mesh:NoExp]) OR “Stress Disorders, Post-Traumatic”[Mesh]) OR “Stress Disorders, Traumatic, Acute”[Mesh]) OR “Anti-Anxiety Agents”[Mesh]) OR “Anti-Anxiety Agents” [Pharmacological Action])) AND ((((((“Animal Experimentation”[Mesh]) OR “Models, Animal”[Mesh:NoExp]) OR “Disease Models, Animal”[Mesh:NoExp]) OR “Animals”[Mesh]) OR “Behavior, Animal”[Mesh]) OR “Animals, Laboratory”[Mesh])). “Other Animals” filter was applied in the research to exclude studies with humans.

Two independent researchers reviewed the articles at one or more of four levels of detail (title, abstract, full text and a detailed review of experimental design) to determine their eligibility. The inclusion criteria adopted were: (1) administration of ketamine (any dose, route and schedule of treatment) as an intervention group and an appropriated control group; (2) primary behavior data from a test related to anxiety; (3) detailed outcome data (mean, standard error or standard deviation, and sample sizes of both control and intervention groups). Studies that administered ketamine at drinking water or that administered a combination of drugs (including ketamine) to the subjects were excluded.

Regarding item 2 (primary behavior data from a test related to anxiety), we considered all behavioral tests that measure any kind of trace or state anxiety or fear elicited by aversive stimuli (Bourin, 2015; Cryan and Sweeney, 2011; Lezak et al., 2017). For example, although there is no consensus that the open field test is an animal model of anxiety, many researchers consider that proportion of time or entries in the center of the open field would reflect trace anxiety (Harro, 2018; Prut et al., 2003). Thus, of the studies that evaluated ketamine effects on the open field, we selected only those studies that reported exploration of the center of the open field.

The selected articles were transferred electronically to Mendeley software (version 1.17.13). We collected the following data from each of the included studies: authors, year of publication, animal details (specie, gender and strain), treatment regimen of ketamine, type of ketamine (racemic mixture, S-ketamine or R-ketamine), test of anxiety/fear and behavioral outcome. In the studies that did not describe the type of ketamine, but did mention the manufacturer of ketamine, we searched the grey literature (google search) to identify the type of ketamine based on manufacturer information. We performed the review of the literature to include articles published between January, 1972 and June, 2018.

### 2.2 Behavioral Analysis

#### 2.2.1 Animals

A total of 164 naïve male adult Wistar rats were obtained from the breeding colony of Federal University of Espirito Santo and were allowed to acclimatize in our laboratory for 3 weeks before starting the tests. They were housed in groups of 4 in Plexiglas cointainers (49 x 34 x 26 cm), under 12 h/12 h at light/dark cycle (lights on at 6:30 am) and in a temperature-controlled room (22-24 °C), with free access to tap water and standard chow diet (Nuvilab CR1, Quimtia S.A., Brazil). Cages were lined with wood shavings and replaced by clean cages 3 times/week. No enrichment material was used inside the cages. At the moment of the tests, rats were weighing 280 to 350 g and aging 12 to 14 weeks. The animals were randomly assigned in independent groups to the different treatment conditions and each rat was tested only once. All behavioral experiments were conducted between 12:00 and 6:00 pm. The experimental procedures were conducted in accordance with the National Council for Control of Animal Experimentation (CONCEA, Brazil) and with EU Directive 2010/63/EU for animal experiments. Protocols were approved by the local Committee for the Ethical Use of Animals in scientific research (CEUA, 048/2014).

#### 2.2.2 Drugs

The following drugs were used: ketamine hydrochloride, racemic mixture (Syntec do Brasil, Cotia, SP, Brazil), a non-competitive NMDA antagonist receptor, was administered by intraperitoneal (i.p.) route at sub-anesthetic (10 and 30 mg/kg) and anesthetic doses (80 mg/kg; doses based on Harkin et al., 2003 and pilot studies in our laboratory); MK-801, also known as dizocilpine (Sigma-Aldrich, St. Louis, MO, USA), a non-competitive NMDA antagonist receptor, was administered by i.p. injection at doses of 0.05 and 0.1 mg/kg (Maeng et al., 2008). All the drugs were freshly dissolved in NaCl 0.9 % (saline). We confirmed that ketamine 80 mg/kg induced anesthesia by checking the lost of tail pinch reflex and pedal withdrawal reflex in the treated animals. Animals that received the sub-anesthetic doses of ketamine did not exhibit sedation or severe ataxia after administration. The animals from control groups received saline 1 ml/kg by i.p. route.

#### 2.2.3 Behavioral Tests

##### 2.2.3.1 Elevated T-maze (ETM)

The ETM was derived from the elevated plus maze and was developed by Graeff and colleagues (Zangrossi and Graeff, 1997) to evaluate, in the same apparatus, conflict-evoked and fear behaviors. Accordingly, the differences in the experimental setups between the EPM and ETM were mostly procedural. The apparatus consists of 3 arms of equal dimension (50 cm × 10 cm) that were elevated from the floor 50-cm. One of these arms is surrounded by 40-cm high walls and perpendicular to 2 open arms. Three days before the test, rats were handled once a day during 5 minutes to reduce the aversion to the experimenter. Twenty-four hours before the test, the animals were subjected to 30-min confinement in an open arm. This procedure facilitates escape by reducing exploration of the open arms (Teixeira et al., 2000). On the next day, the acquisition of inhibitory avoidance was assessed by placing the rat at the end of the closed arm and measuring the time that the rat took to leave the closed arm in 3 consecutive trials (baseline, inhibitory avoidance 1 and 2). Escape response was assessed by placing the animal on the extremity of the previously explored open arm and measuring the latency that the animal took to leave this arm in three successive trials (escape 1, escape 2 and escape 3). A cut off time for inhibitory avoidance and escape was 300 s. The experimenter was blind to the pharmacological treatment.

##### 2.2.3.2 Open field test (OFT)

The arena (open field) consists of a wooden box with 1 m^2^ area and 30-cm high walls. Immediately after the ETM test, the rats were placed in the center of the arena to measure locomotor activity for 5 min. The arena was cleaned with a water-alcohol solution (10%) after each animal was tested. The tests were videorecorded with a webcam that was connected to a computer. Total distance traveled (m) was analyzed by the Anymaze 4.98 path-tracking software (Stoelting, Wood Dale, IL, USA).

### 2.3 Experimental Design

#### 2.3.1 Experiment 1 – Evaluation of acute effect of systemic administration of ketamine and MK-801 in the ETM and OFT

This experiment was designed to evaluate acute effects of both ketamine and MK-801 in the ETM. Firstly, the animals were submitted to a pre-exposure to one of the open arms of the maze. On the test day, the animals received the injection of ketamine at doses of 10 and 30 mg/kg (sub-anesthetic doses) or saline i.p. The test was performed 45 min after injection. An independent group of animals received one injection of ketamine 80 mg/kg (anesthetic dose) or saline 2 hours before the test, to ensure that the animals would not be sedated at the moment of the tests (based in a pilot study). The last independent group of animals was treated with MK-801 at doses of 0.05 and 0.1 mg/kg or saline 45 min before the test. All animals were tested in the ETM and immediately after submitted to the OFT to evaluate their spontaneous locomotor activity.

#### 2.3.2 Experiment 2 - Evaluation of sustained effect of systemic administration of ketamine and MK-801 in the ETM and OFT

In this experiment, the animals were treated with ketamine (10, 30 and 80 mg/kg), MK-801 (0.05 and 0.1 mg/kg) or saline i.p. 24 hours before the test to evaluate possible sustained effects in the ETM and in the OFT.

### 2.4 Statistical analysis

Data analyses were performed using Statistical Package for the Social Sciences SPSS 20.0 software (IBM SPSS Statistics^®^, Chicago, IL, USA). The homogeneity of variances was assessed using Levene test. When Levene’s test showed that data were not homocedastic, data were log-transformed previously to analyses. Data from ETM (inhibitory avoidance and escape) were analyzed using a two-way repeated measure analysis of variance (ANOVA), with treatment as the independent factor and trials (baseline, avoidance 1–2, and escape 1–3) as the dependent factors. Significant effects of the independent factor or the interaction between the independent and dependent factors were followed by Dunnett *post hoc* test. Data from the OFT were analyzed by one-way ANOVA followed by Dunnett *post hoc* test when appropriate. Data from the OFT of experiment that evaluated the anesthetic dose of ketamine were analyzed using unpaired Student’s t-test. The significance level was set at p < 0.05. Data were expressed as mean ± SEM.

## 3 Results

### 3.1 Literature Review

After applying the MeSH® words previously described, we found 2038 studies in our literature search. Thus, we applied the filter “Other Animals” to refine our search and found 795 studies. After a check against inclusion and exclusion criteria, we initially selected 51 studies that evaluated ketamine effects in animal tests of anxiety/fear. We excluded 3 studies that evaluated ketamine effects in animals chronically exposed to ethanol. We also excluded 1 study that administered ketamine in drinking water and 2 studies that did not exhibit outcome data (mean and S.E.M.).

Table 1 summarizes data extracted from the 45 selected studies. The reporting of details was variable across studies. Fifty five percent of all selected studies were performed in rats, 37 % were performed in mice and only 13 % included females in their evaluation. The vast majority of studies tested ketamine in adult animals. However, in 8 studies, the age of animals was not reported. These studies described the weight of the animals, from which age cannot be properly identified.

**Table 1.**
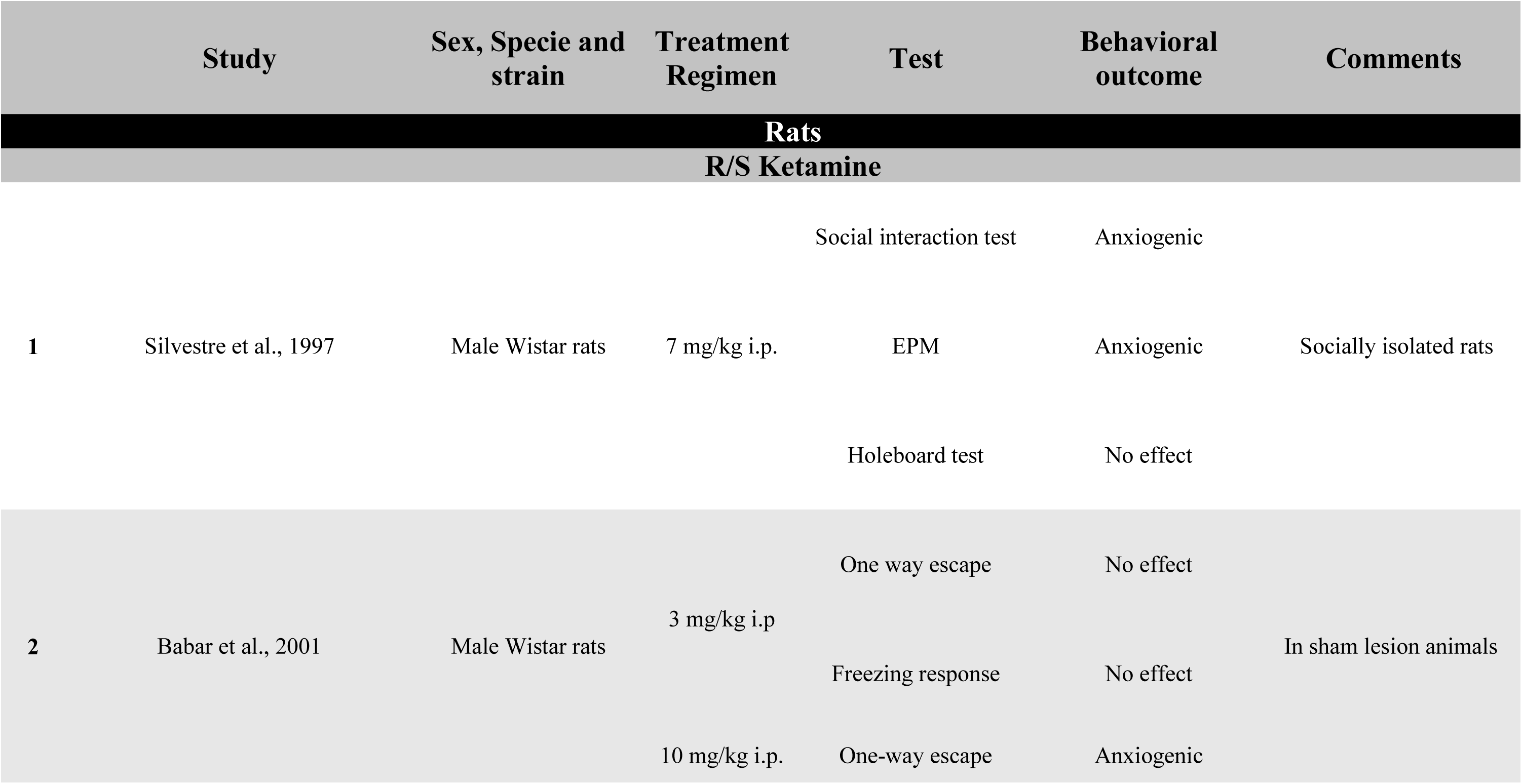
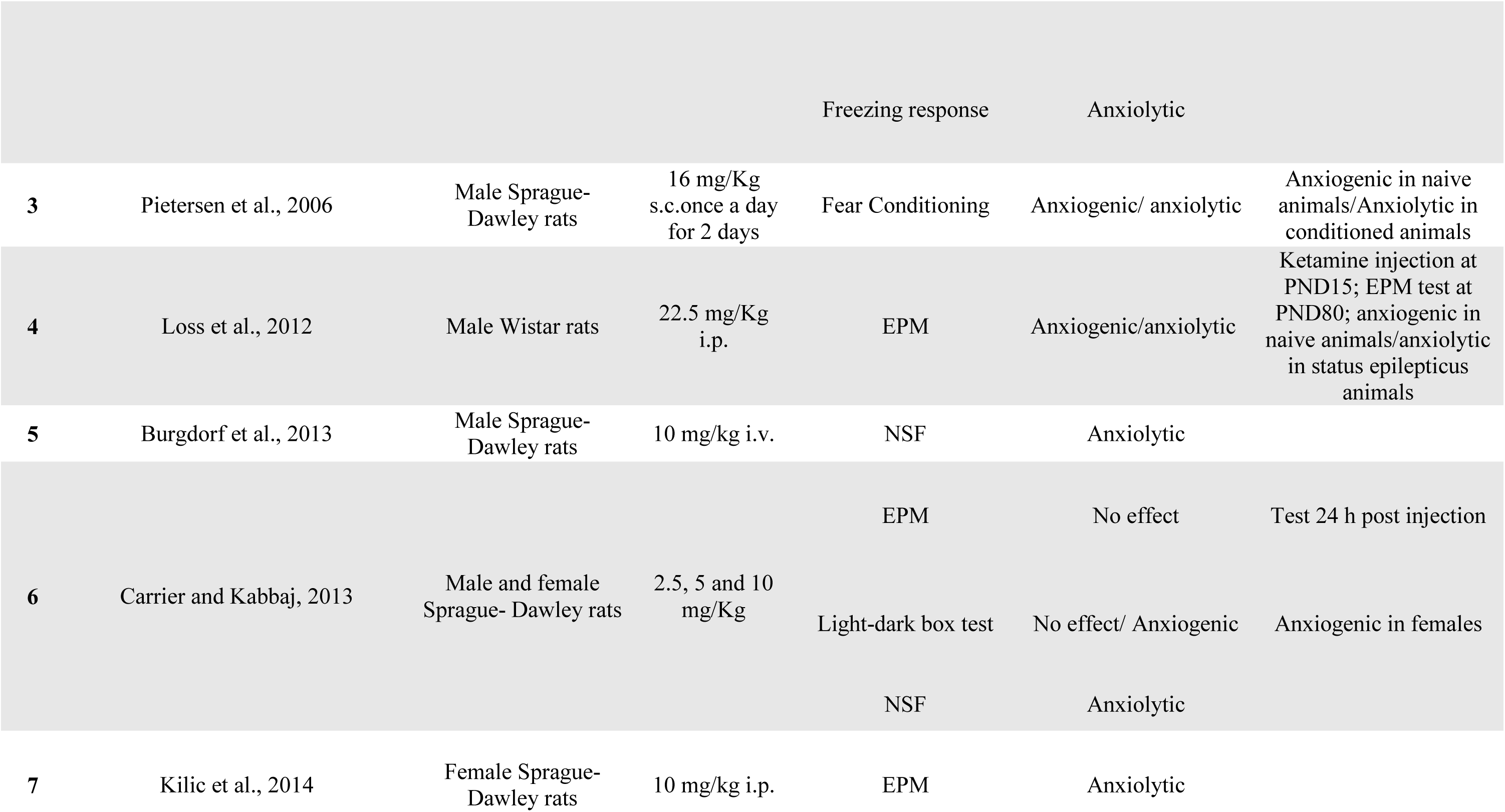
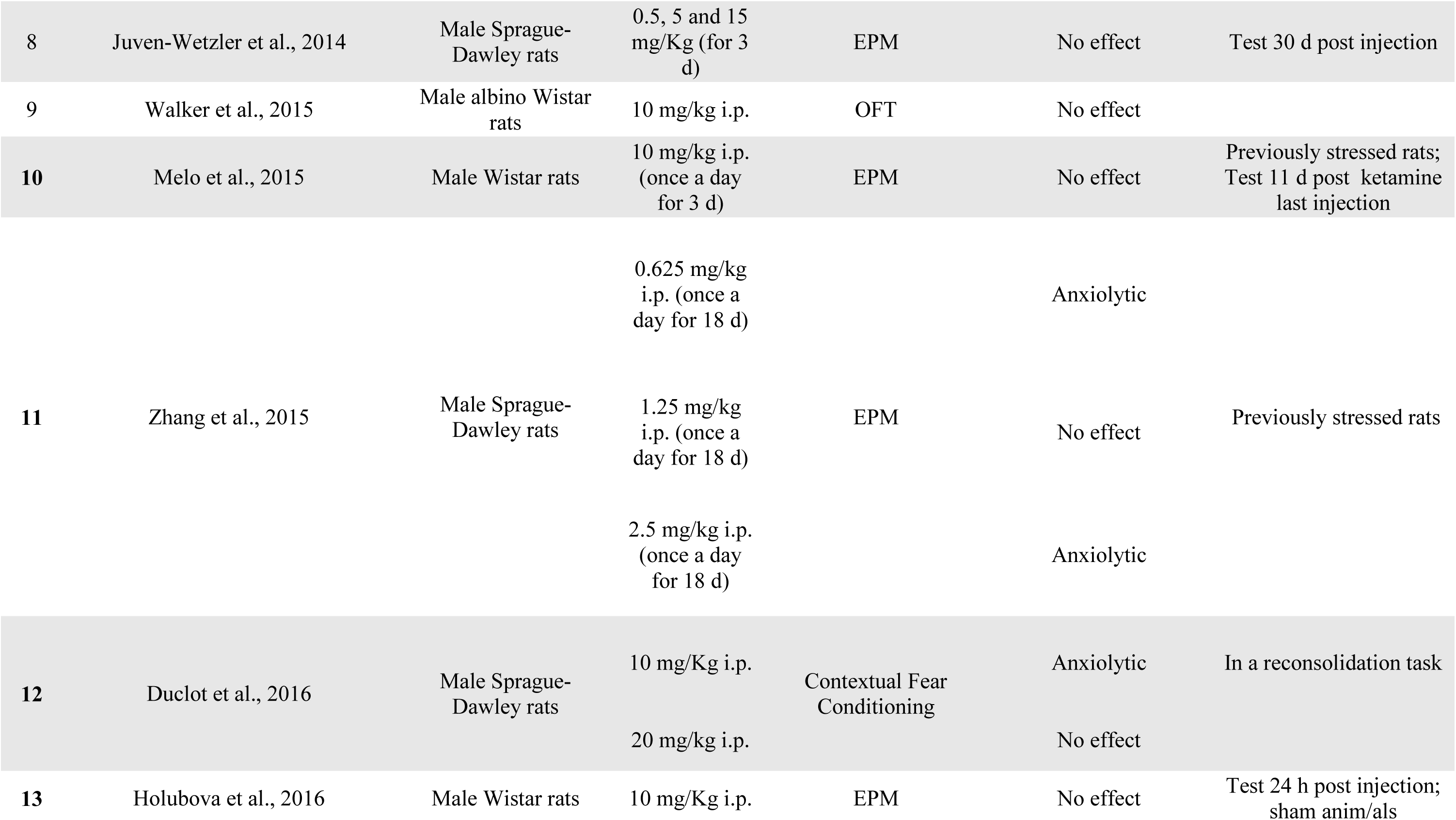
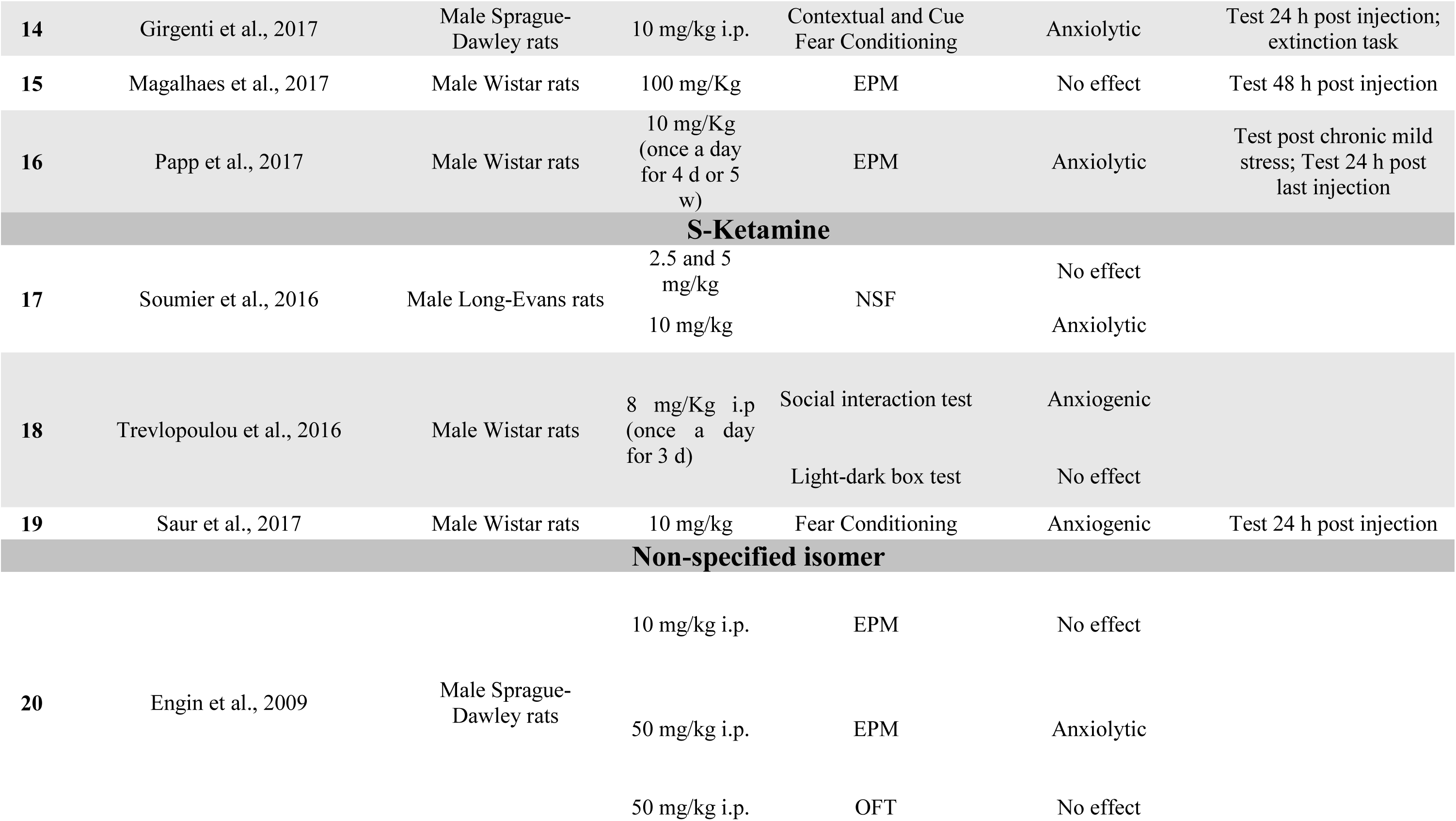
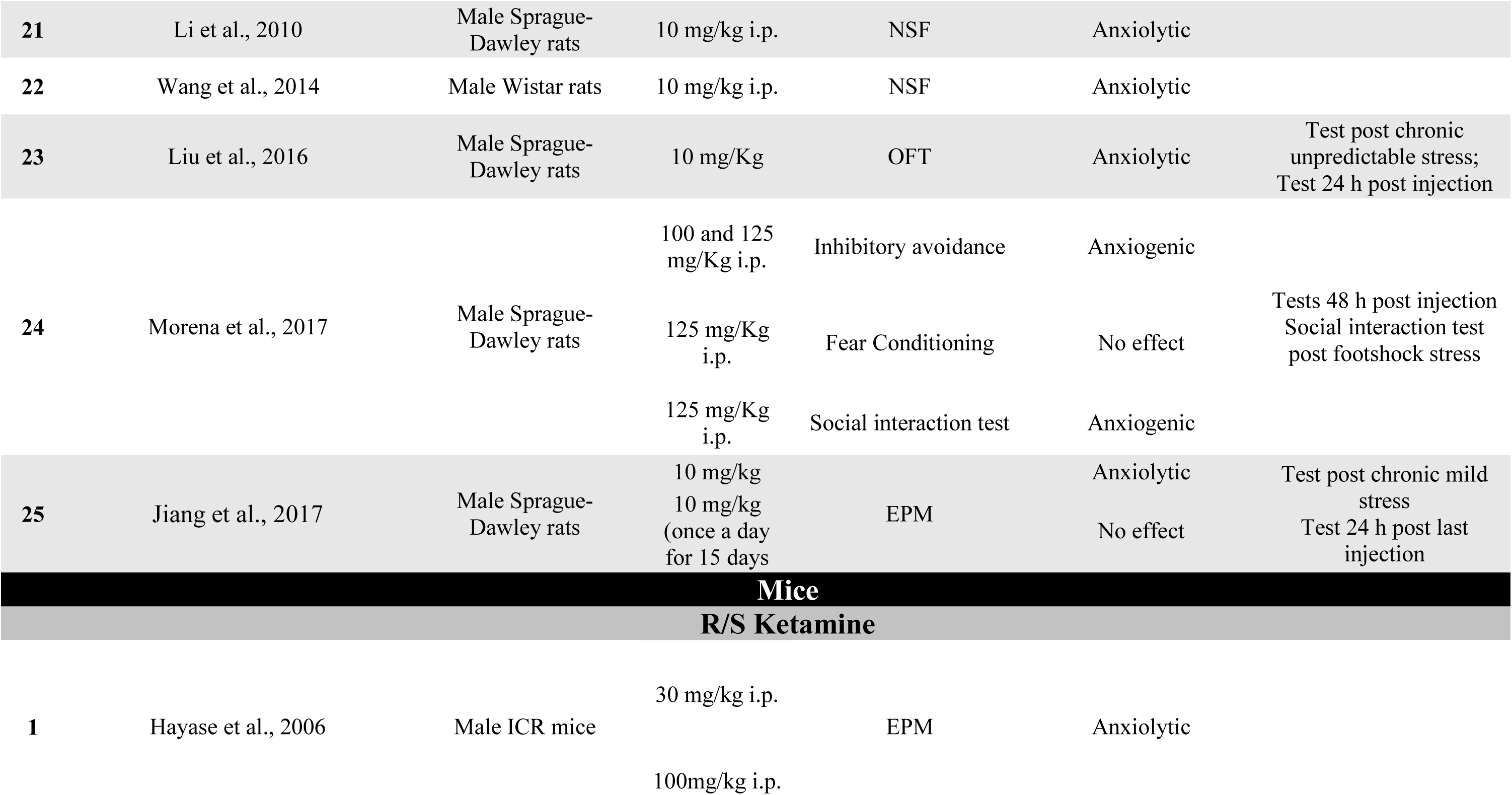
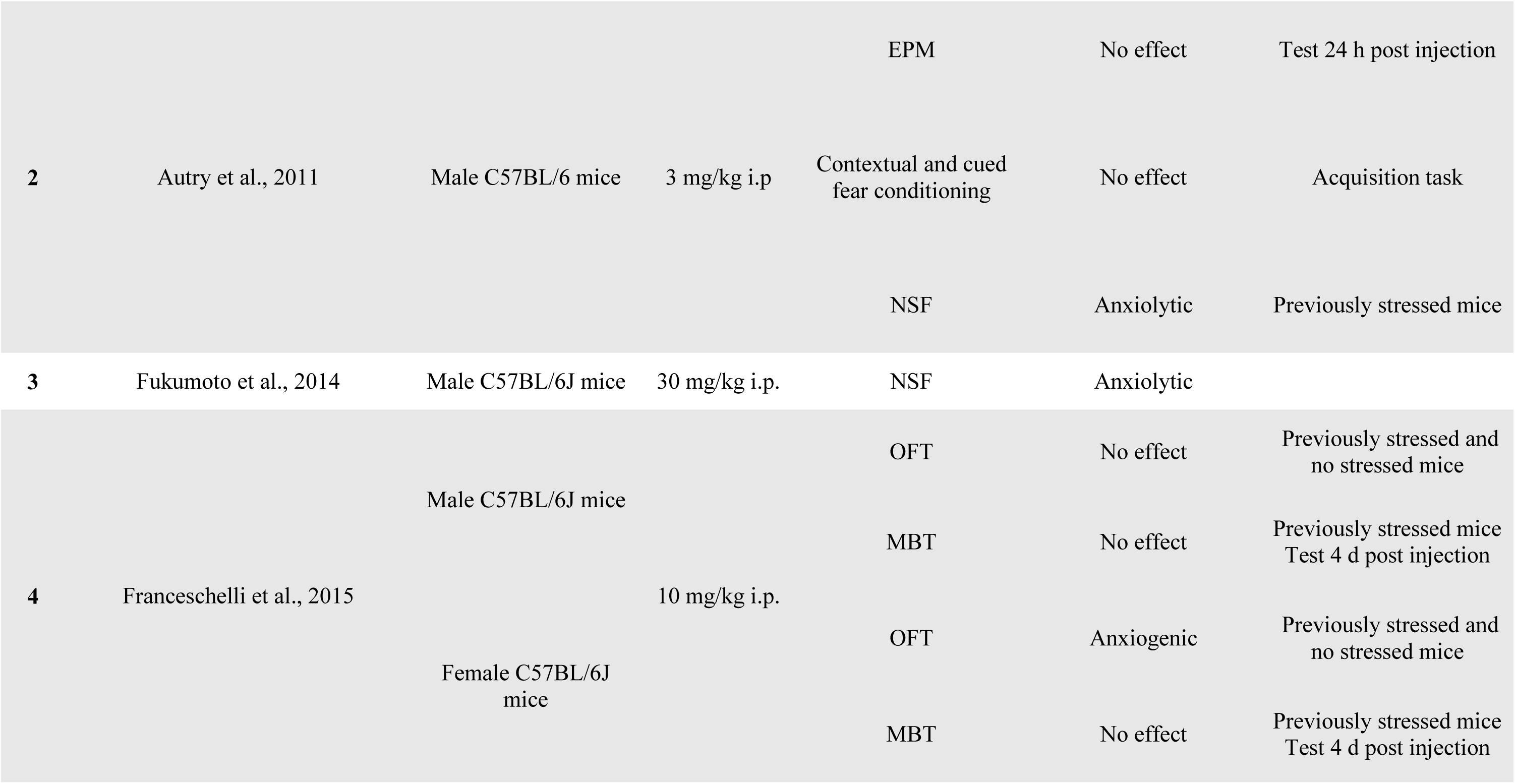
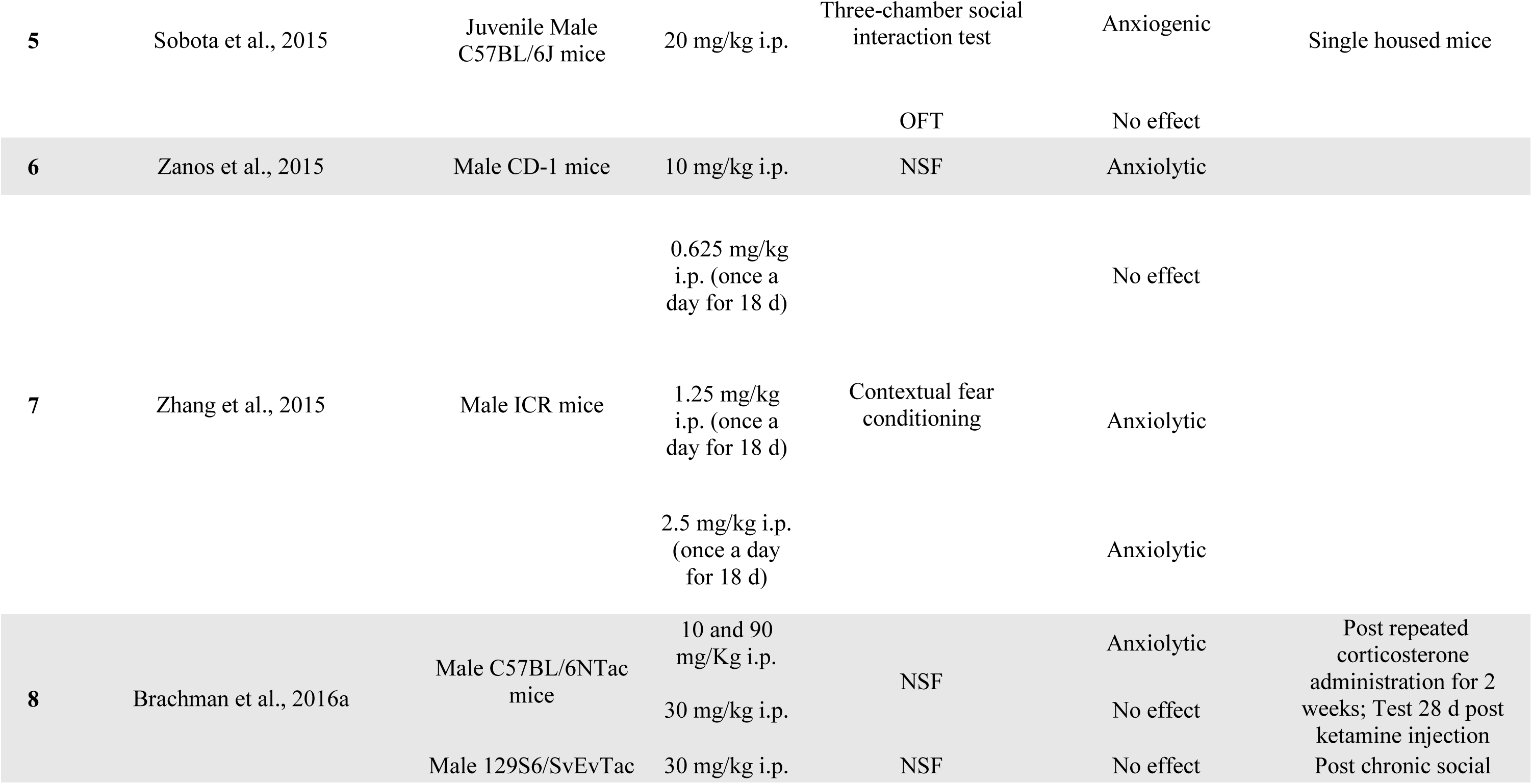
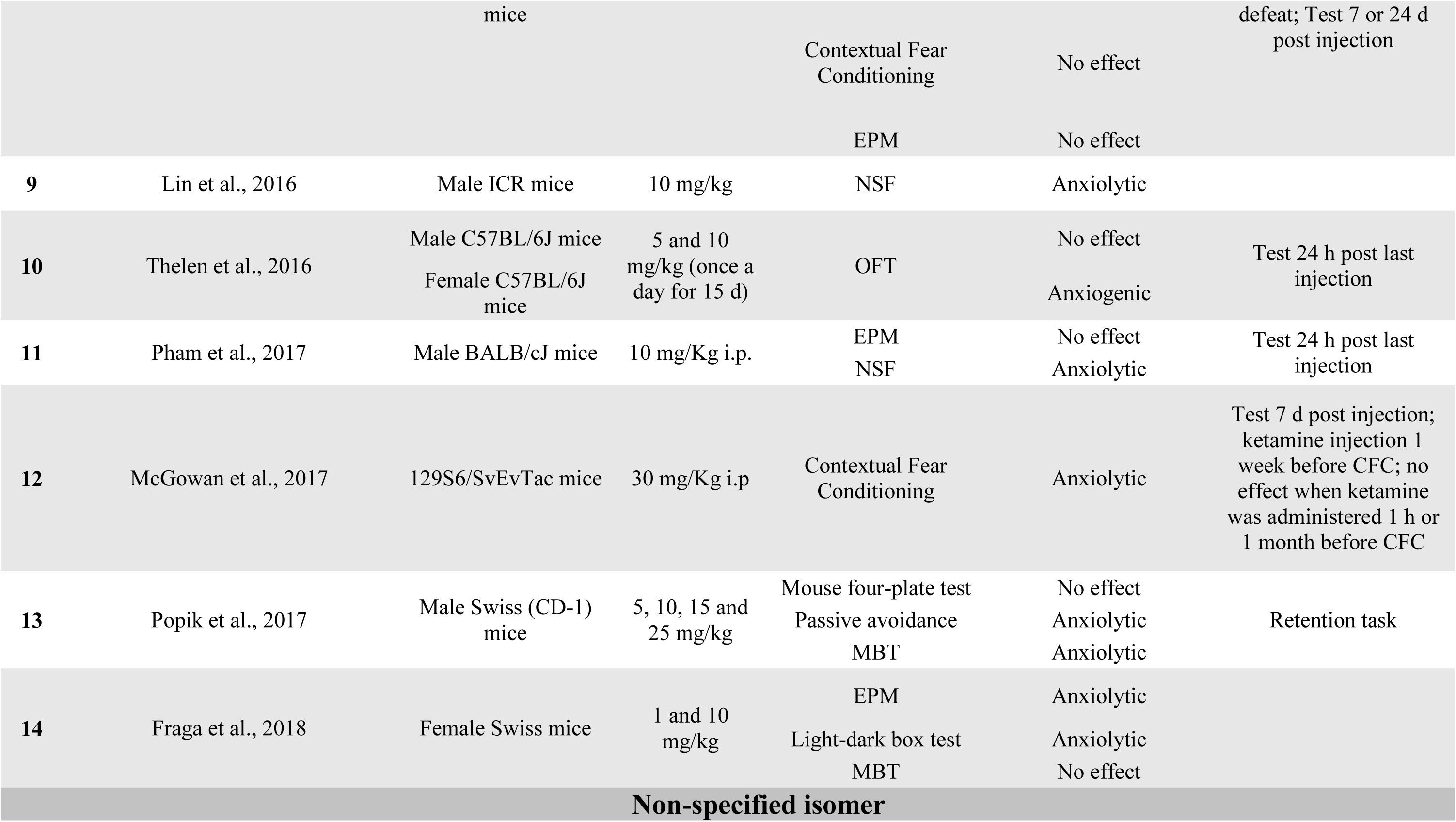
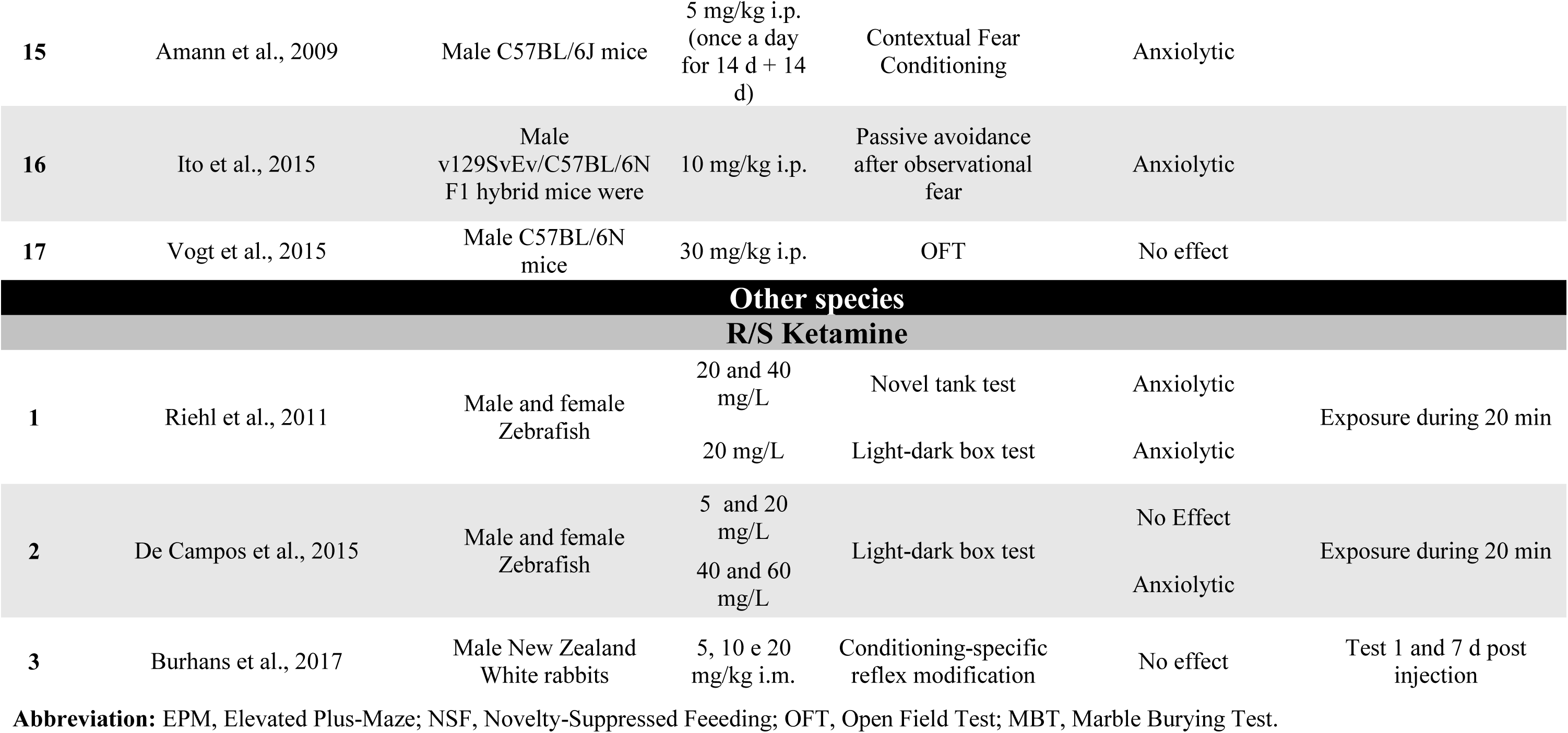
Effects of ketamine on anxiety/fear-related behaviors.

The majority of studies did not report directly which type of ketamine (racemic mixture or isomer) was used. Thus, we searched the grey literature (google search) to identify the type of ketamine based on manufacturer information. In 9 studies we were still unable to identify the type of ketamine. Of the total of selected studies, only 4 evaluated the effects of S-ketamine. No studies evaluated the effects of R-ketamine on anxiety/fear-related behaviors. Only 4 experiments evaluated the effects of anesthetic doses of ketamine.

Most of the selected studies assessed anxiety/fear-related behaviors using the EPM, NSF and fear conditioning. No selected studies evaluated the effect of ketamine in animal tests related to PD. Regarding the outcome of studies performed in rats, 14 experiments detected an anxiolytic-like effect of ketamine, 9 experiments detected an anxiogenic-like effect and 19 described null effect of ketamine. From the experiments in which ketamine had null effect (total number 19), 8 tested doses of ketamine under 10 mg/kg (potentially sub-effective doses) and 6 evaluated anxiety-related behaviors at least 24 h after ketamine administration.

In experiments performed in mice, the pattern of results was similar. Twelve experiments detected an anxiolytic-like effect of ketamine, 2 experiments detected an anxiogenic-like effect and 10 described null effect of ketamine. From the experiments describing that ketamine had null effect (total number 10), 6 tested doses of ketamine under 10 mg/kg (potentially sub-effective doses) and 3 evaluated anxiety-related behaviors at least 24 h after ketamine administration.

### 3.2 Behavioral Analysis

#### 3.2.1 Experiment 1 - Evaluation of acute effect of systemic administration of ketamine and MK-801 in the ETM and OFT

The sub-anesthetic doses of ketamine (10 and 30 mg/kg) did not affect the inhibitory avoidance performance [trial effect: F_(2, 76)_= 33.937; p< 0.001; treatment: F_(2, 38)_= 0.289; p= 0.751; interaction between treatment and trial: F_(4, 76)_= 1.373; p= 0.251; Fig. 1A] and escape performance [Trial effect: F_(1.569, 76)_= 0.852; p= 0.370; treatment: F_(2, 38)_= 0.119; p= 0.888; interaction between treatment and trial: F_(3.138, 76)_= 1.599; p= 0.197 (Fig. 1B) in the ETM. The sub-anesthetic doses of ketamine also did not impair locomotor activity in the OFT (F_(2, 38)_= 0.23; p= 0.795; Table 2).

**Figure 1.**
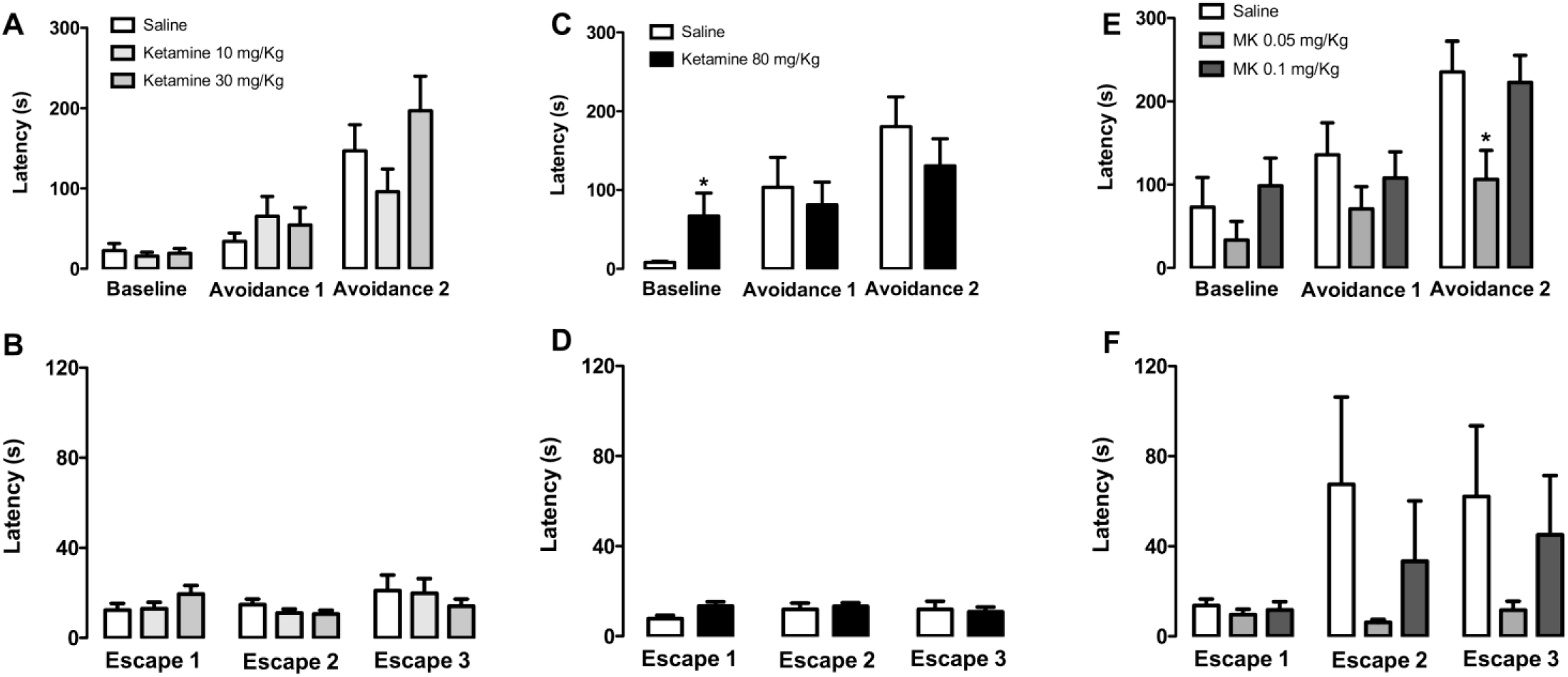
Acute effects of systemic administration (i.p.) of ketamine and MK-801 on behavioral tasks in the elevated T-maze. **A** and **B** represent inhibitory avoidance and escape, respectively, from rats treated with ketamine 10 mg/Kg (n=14), 30 mg/Kg (n=12) or saline (control group; n= 15) 45 min before the test; **C** and **D** represent inhibitory avoidance and escape, respectively, from rats treated with ketamine 80 mg/Kg (n=16) or saline (n= 13) 2 h before the test; **E** and **F** represent inhibitory avoidance and escape, respectively, from rats treated with MK-801 0.05 mg/Kg (n= 13), 0.1 mg/Kg (n= 11) or saline (n= 10) 45 min before the test. *p < 0.05 compared to control group in the same trial (two-way ANOVA followed by Dunnett’s *post hoc* test). The data are expressed as mean ± SEM.

**Table 2.** Effects of ketamine and MK-801 on the distance traveled (means ± S.E.M) by rats in the open-field.

| **Treatment** | **Total Distance Traveled**<br>**(m)** |
| --- | --- |
| **Sub-anesthetic doses of ketamine administered 45 min before the test** |  |
| Saline | 17.30 ± 3.76 |
| Ketamine 10 mg/kg | 12.98 ± 1.71 |
| Ketamine 30 mg/kg | 17.44 ± 2.24 |
| **Anesthetic dose of ketamine administered 2 h before the test** |  |
| Saline | 18.43 ± 1.94 |
| Ketamine 80 mg/kg | 24.21 ± 3.13 |
| **MK-801 administered 45 min before the test** |  |
| Saline | 11.44 ± 2.46 |
| MK-801 0.05 mg/kg | 14.73 ± 18.8 |
| Mk-801 0.1 mg/kg | 18.14 ± 3.31 |
| **Ketamine administered 24 hours before the test** |  |
| Saline | 19.11 ± 2.18 |
| Ketamine 10 mg/kg | 18.31 ± 2.01 |
| Ketamine 30 mg/kg | 19.03 ± 1.87 |
| Ketamine 80 mg/kg | 14.96 ± 1.23 |
| **MK-801 administered 24 hours before the test** |  |
| Saline | 16.62 ± 13.8 |
| MK-801 0.05 mg/kg | 19.78 ± 1.73 |

The anesthetic dose of ketamine (80 mg/kg) changed inhibitory avoidance performance in the ETM [trial effect: F_(2, 54)_= 15.434; p< 0.001; treatment: F_(1, 27)_= 0.015; p= 0.903; interaction between treatment and trial: F_(2, 54)_= 3.534; p= 0.036; Fig. 1C]. Specifically, the anesthetic dose of ketamine increased baseline avoidance (t_(27)_= -2.020; p= 0.062) but not avoidance 1 (t_(27)_= 0.482; p= 0.633) and avoidance 2 (t_(27)_= 0.974; p= 0.339). Ketamine 80 mg/kg did not affect the escape behavior [trial effect: F_(2, 54)_= 0.527; p= 0.593; treatment: F_(1, 27)_= 0.702; p= 0.410; interaction between treatment and trial: F_(2, 54)_= 1.462; p= 0.241;; Fig. 1D] in the ETM nor the total distance traveled in the OFT (t_(27)_= -1.482; p= 0.150; Table 2).

We also evaluated the effects of another non-competitive NMDA receptor antagonist, MK-801, in the ETM. Systemic administration of MK-801 changed inhibitory avoidance performance in the ETM [trial effect: F_(2, 62)_= 15.491; p< 0.001; treatment: F_(2, 31)_= 4.113; p= 0.026; interaction between treatment and trial: F _(4, 62)_= 0.778; p= 0.543). Dunnett’s *post hoc* test showed that the lowest dose of MK-801 (0.05 mg/kg) impaired inhibitory avoidance compared to the control group (p < 0.05; Fig. 1E). MK-801 did not impair the escape performance [trial effect: F_(1.242, 38.505)_= 4.046; p< 0.05; treatment: F_(2, 31)_= 1.494; p= 0.240; interaction between treatment and trial: F_(2.484, 38.505)_= 1.456; p= 0.245; Fig. 1.F] in the ETM nor the locomotor activity in the OFT (F_(2, 31)_= 1.581; p= 0.222; Table 2).

#### 3.2.2 Experiment 2 - Evaluation of sustained effect of systemic administration of ketamine and MK-801 in the ETM and OFT

Administration of ketamine 24 h before the ETM did not affect the inhibitory avoidance [trial effect: F_(2, 72)_= 28.867; p< 0.001; treatment: F_(3, 36)_= 0.501; p= 0.684; interaction between treatment and trial: F_(6, 72)_= 2.011; p= 0.075; Fig. 2A] and the escape performance [trial effect: F_(2, 72)_= 5.907; p< 0.05; treatment: F_(3, 36)_= 0.944; p= 0.430; interaction between treatment and trial: F_(6, 72)_= 1.202; p= 0.315; Fig. 2B]. Also, ketamine, at all tested doses, did not impair the locomotor activity in the OFT (F_(3, 36)_= 1.243; p= 0.309; Table 2).

**Figure 2.**
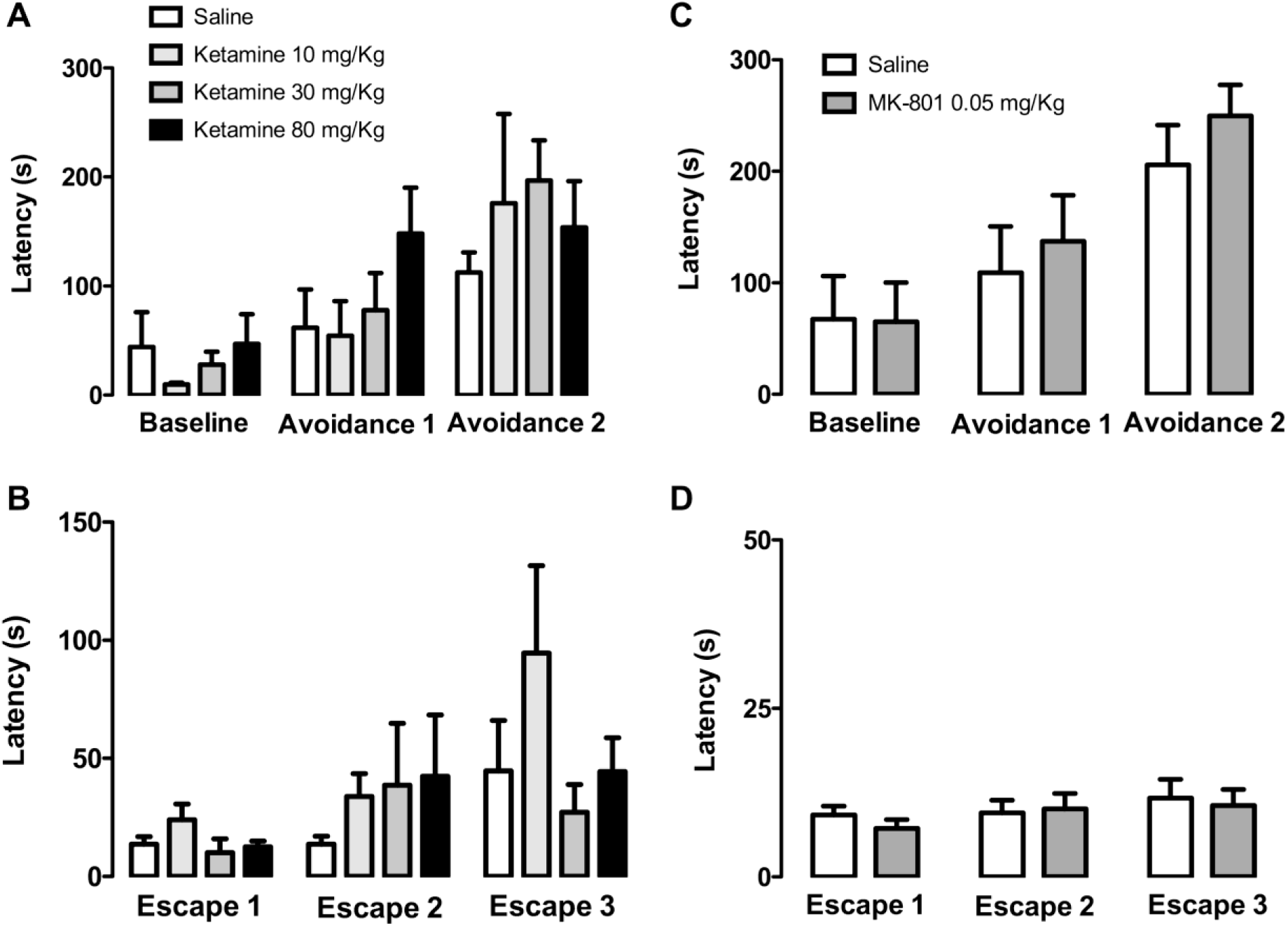
Evaluation of putative sustained effects of systemic administration (i.p.) of ketamine and MK-801 24 hours before the behavioral tasks in the elevated T-maze. **A** and **B** represent inhibitory avoidance and escape, respectively, from rats treated with ketamine 10 (n=9), 30 (n=11) and 80 mg/Kg (n= 11) or saline (control group; n= 9) 24 hours before the test; **C** and **D** represent inhibitory avoidance and escape, respectively, from rats treated with MK-801 0.05 mg/Kg (n= 11) or saline (n= 10) 24 hours before the test. The data are expressed as mean ± SEM.

As a single injection of MK-801 0.05 mg/kg acutely impaired inhibitory avoidance in the ETM, we tested if this dose of MK-801 would induce a sustained anxiolytic-like effect. We showed that prior administration of MK-801 0.05 mg/Kg (24 h before the test) did not affect the inhibitory performance [trial effect: F_(2, 38)_= 17.961; p< 0.001; treatment: F_(1,19)_= 0.316; p= 0.581; interaction between trial and treatment: F_(2,38)_= 0.366; p= 0.696; Fig. 2C] and the escape performance [trial effect: F_(2, 38)_= 1.845; p= 0.172; treatment: F_(1,19)_= 0.123; p= 0.730; interaction between trial and treatment: F_(2,38)_= 0.362; p= 0.699; Fig. 2D] in the ETM test. Prior administration of MK-801 also did not alter the locomotor activity in the OFT (t_(19)_= - 1.406; p= 0.176, Table 2).

## 4 Discussion

### 4.1. Literature review

To the best of our knowledge, the present study was the first review addressing ketamine effects on preclinical anxiety/fear tests. Thus, we chose to perform a very comprehensive search which resulted in the selection of studies substantially heterogeneity. There appears to be no pattern in the outcome of the experiments that evaluated the effects of ketamine on anxiety/fear-related behaviors.

Among the factors that may result in variability of ketamine effects, it stands out the differences in the methods, such as: the type and intensity of anxiety evoked by the paradigm and previous interventions; specie/strain studied, including age and sex; type of ketamine used; great range of ketamine doses and variable schedule of treatment; variable interval between ketamine administration and behavioral evaluation.

The Research Domain Criteria (RDoC), an initiative of the National Institute of Mental Health (NIMH), proposed categories to investigate mental health that includes negative affect, positive affect and cognition (Carcone and Ruocco, 2017; Insel et al., 2010). Each of these domains can be tested by different behavioral paradigms. Most anxiety paradigms used in the studies revised herein are linked to negative affect and involve acute threat (fear), potential threat (anxiety) and sustained threat. Moreover, a common prior intervention adopted in studies investigating anxiety/fear-related behaviors is to submit the subjects to stress conditions. Finally, the sustained antidepressant-like effect of ketamine may be influenced somewhat by stress condition (Polis et al., 2019). Therefore, we compared the main effects induced by ketamine on the potential threat (anxiety) models (EPM, OFT and LDT) performed in naive versus previously stressed animals. We found a mix of outcomes (anxiolytic, anxiogenic and no effect) that was very similar in both types of conditions (naive versus stressed animals). Given that, besides the type of anxiety paradigm used in the revised studies, other factors may have contributed to the outcome of ketamine on anxiety/fear-related behaviors. As mentioned in the results section, there are a great variability of dose, number of administration and even interval between administration of ketamine and test exposure.

Regarding published studies involving acute or sustained threat (fear), about half of them reported an anxiolytic-like effect of ketamine in behavioral responses such as freezing. In contrast, all the 3 experiments that evaluated social interaction as a consequence of acute or sustained stress reported an anxiogenic-like effect of ketamine. Noteworthy, most of these studies did not evaluate ketamine effects considering resilient versus stress-vulnerable animals. As proposed by Musazzi and coworkers (Musazzi et al., 2018), investigate ketamine effects on resilient versus stress-vulnerable animals could increase accuracy of results.

In general, paradigms of acute or sustained threat have been related to PTSD (Frandreau and Toth, 2017). There are few studies that evaluated ketamine effects on patients diagnosed with PTSD or with symptoms of PTSD (Banov et al., 2019; Liriano et al., 2019). Although some of these studies have shown that ketamine may be an option for PTSD treatment (Albott et al., 2018; Feder et al., 2014), other studies described possible side effects of ketamine that could worsen PTSD symptoms (Schönenberg et al., 2008, 2005). Of note, this last study mentioned found that accident victims treated with S-ketamine exhibited greater symptoms of dissociation, re-experiencing and avoidance than those treated with racemic ketamine (Schönenberg et al., 2005). On the other hand, depressed patients with a history of PTSD who received a single racemic ketamine infusion did not experience a clinically significant increase in dissociative or anxiety symptoms (Zeng et al., 2013). These conflicting and pending results highlight the need for further studies comparing S-ketamine and R-ketamine in preclinical protocols of acute threat as well as in patients diagnosed with PTSD with and without comorbidities, such as depression.

In contrast, from the 12 experiments that evaluated ketamine effects on NSF, 11 reported an anxiolytic-like effect of ketamine and only 1 report no effect. The NSF is based on the inhibition of feeding behavior generally caused by a novel environment (Blasco-Serra et al., 2017). The NSF has been considered an animal model of anxiety since it measures a conflict between the anxiogenic environment and hunger-induced behavior, which is sensitive to anxiolytic drugs (Cryan and Sweeney, 2011; Dulawa and Hen, 2005; Merali et al., 2003; Shephard and Broadhurst, 1982). On the other hand, some authors suggest that NSF could be an animal model of depression by assessing anhedonia and detecting the effects of chronic, but not acute treatment with antidepressant drugs (Dulawa and Hen, 2005; Nestler and Hyman, 2010; Powell et al., 2012). Thus, one possibility is that the test assesses antidepressant drug effects by detecting their anxiolytic effects (Ramaker and Dulawa, 2017). Within this context, many of the selected studies by the present review reported the effect of ketamine on NSF as a fast-acting antidepressant-like effect. Of note, some of the selected studies reported that ketamine had no effect on distinct anxiety-like behaviors (Autry et al., 2011; Brachman et al., 2016; Carrier and Kabbaj, 2013; Pham et al., 2017) while others reported that ketamine had an antidepressant-like effect in distinct depression-like behaviors (Li et al., 2010; Lin et al., 2016; Soumier et al., 2016; Zanos et al., 2015). In this way, the total data set on the NSF is most consistent with an antidepressant-like effect of ketamine rather than an anxiolytic-like effect.

The direction of ketamine effects on anxiety/fear-related behaviors might also be influenced by the so-called publication bias. There is some evidence suggesting that negative results are underreported/unpublished (Driessen et al., 2015; Mlinarić et al., 2017). In fact, the risk of publication bias in both, pre-clinical (Mohammad et al., 2016) and clinical studies (Turner et al., 2008), have already been identified. Therefore, there is much space to improve pre-clinical research and reduce the risk of bias. For example, a good practice is to register, in advance, the protocols and experimental design of projects in online repositories such as www.preclinicaltrials.eu, to reduce risk of bias (Jansen et al., 2014; Tillmann, 2017).

The vast majority of the revised articles did not consider gender differences in their experiments. Although is not completely understood whether ketamine has sex-specific effects (Chang et al., 2018; Herzog et al., 2019), some evidence suggests more pronounced biochemical and behavioral effects of ketamine in females (Carrier and Kabbaj, 2013; Dossat et al., 2018; Franceschelli et al., 2015; Guarraci et al., 2018; Saland and Kabbaj, 2018; Thelen et al., 2019, 2016). It is noteworthy that women are twice more likely than men to experience an anxiety disorder over their lifespan (Bandelow and Michaelis, 2015; Craske et al., 2017; Hantsoo and Epperson, 2017; Remes et al., 2016). Altogether, these data highlight the importance of filling this gap, which means to produce further evaluation of ketamine effects on anxiety/fear-related behavior in females.

Regarding the quality of the selected studies, some points deserve to be mentioned. Ketamine is a chiral anesthetic drug that is usually marketed as a racemic mixture containing equal amounts of two enantiomers, S- and R-ketamine. Meanwhile, pharmacokinetic and pharmacodynamic differences between the ketamine enantiomers have been described (Muller et al., 2016). For instance, S-ketamine has about a 2-4-fold greater affinity for the NMDA and opioid receptors, higher anesthetic potency and maybe fewer adverse effects than R-ketamine (Andrade, 2017; Domino, 2010; Kohrs and Durieux, 1998; Muller et al., 2016; White et al., 1980). In constrast, R-ketamine seems to have a longer lasting antidepressant-like effect than S-ketamine possibly mediated by different molecular mechanisms (Fukumoto et al., 2017; Yang et al., 2018, 2015; Zhang et al., 2014). Surprisingly, most of the selected studies in this review were inaccurate in identifying the type of ketamine that was used and some of them did not even describe the manufacturer. Additionally, in our review, in 3 of the studies that specifically investigated the effects of S-ketamine on anxiety/fear-related behaviors, 2 identified an anxiogenic-like effect of the drug. From this, it is important that future studies reduce the risk of bias by clearly describing the type of ketamine used and its manufacturer. Also, future studies should address specifically the effects of different enantiomers of ketamine on anxiety/fear-related behaviors.

Another issue about the quality of the selected studies is the absence of detailed information about the interval between ketamine administration and behavioral evaluation. This observation is in line with previous evidence suggesting that the majority of publications fail to include key information (Avey et al., 2016; Kilkenny et al., 2009; Macleod et al., 2015). Inaccurate or incomplete reporting has potential scientific, ethical, and economic implications. Collectively, these data strengthen the importance of following guidelines such as those of PREPARE (Planning Research and Experimental Procedures on Animals: Recommendations for Excellence)(Smith et al., 2018) and ARRIVE (Animals in Research: Reporting In Vivo Experiments) (Kilkenny et al., 2010; Percie et al., 2019) initiative to increase reproducibility and to reduce publication bias in pre-clinical studies.

Finally, it is important to mention some limitations of our review. We limited our search to the electronic database PubMed and to English-language published data. We also did not perform a meta-analysis of the data.

### 4.2. New findings

From the evidence gathered in our review, we identified an important gap in the literature regarding ketamine effects in fear-related behaviors associated to PD. Therefore, we evaluated the effect of ketamine in an animal model related to PD. Overall, we found that: 1) racemic ketamine, at sub-anesthetic and anesthesic doses, did not induce acute and/or sustained anxiolytic/panicolytic-like effects in rats exposed to the ETM; 2) MK-801, another non-competitive NMDA receptor antagonist, induced an acute, but not sustained anxiolytic-like effect in rats exposed to the ETM. The reported effects do not seem to be due to locomotor activity changes, as both drugs, at any administered dose and interval of evaluation, did not influence open field performance. Despite this, the anesthetic dose of ketamine increased baseline latency during inhibitory avoidance, suggesting at least a residual depressor effect.

The lack of ketamine effects on the ETM rats performance, acutely or 24 h after its administration, reinforces the controversial findings reported in previous studies, as described in table 1. For example, anesthetic doses of ketamine induced anxiolytic-like (Hayase et al., 2006), anxiogenic-like (Morena et al., 2017) or no effect (Magalhaes et al., 2017) in different tests of anxiety. Regarding sub-anesthetic doses of ketamine, similar mix of outcomes has been previously reported (as detailed in Table 1). Although the reasons for these discrepant results are unknown, they may be attributed to differences in the behavioral paradigms, animal species and interval between ketamine treatment and test. For instance, the genetic background of a strain is among the several factors that may affect the behavioral responses of the animals to a drug (An et al., 2011; Võikar et al., 2001). Ketamine 30 mg/kg produced an anxiolytic-like effect on male ICR mice (Hayase et al., 2006) and no effect on male 129S6/SvEvTac mice (Brachman et al., 2016) submitted to the EPM. Ketamine 10 mg/kg induced an anxiolytic-like effect in NSF, but did not alter the behavior in the EPM and in the context and cued fear conditioning (Autry et al., 2011). One possible explanation is that different animal models of anxiety can recruit distinct neural substrates and, therefore, produce contradictory results.

We also cannot exclude that ketamine effects on anxiety-like behaviors are influenced by the ketamine enantiomer. As described in the previous section, S- and R-ketamine have different neurochemical and behavioural effects and in the present study, we evaluated a racemic mixture of ketamine on behavioral tests. Thus, we cannot exclude the possibility that an specific enantiomer of ketamine has a potential panicolytic-like effect.

Noteworthy, the lowest tested dose of MK-801, a highly selective NMDA receptor antagonist (Wong et al., 1986), impaired inhibitory avoidance, but not escape behavior, suggesting an acute anxiolytic-like effect. This result is in agreement with results observed in other studies that evaluated the effect of MK-801 in animal models of anxiety (Adamec et al., 1998; Engin et al., 2009; Karcz-Kubicha et al., 1997; Refsgaard et al., 2017). Additionally, the anxiolytic-like effect of MK-801 was not sustained, as it was not detected 24 h later. This absence of sustained effect of MK-801 in the ETM is consistent with the absence of sustained effects in the EPM (Hill et al., 2015) and in the forced swimming test (Autry et al., 2011).

One speculative explanation for the contradictory effects of ketamine and MK-801, both NMDA receptor antagonists, is that ketamine, different from MK-801, is a multimodal-acting drug. Ketamine and its active metabolites can act beyond the NMDA receptor, several other targets such as opioid and serotonergic receptors, calcium, sodium and chloride channels, 5-HT transporter, among others (du Jardin et al., 2016; Kapur and Seeman, 2001; Salat et al., 2015). In this sense, it is possible that ketamine and its metabolites could act on different types of receptors with opposite transduction mechanisms that would prevent the appearance of an anxiolytic-like effect.

Another possibility is a differential modulation of glutamate release by ketamine and MK-801 in the prefrontal cortex. Systemic ketamine administration (10-30 mg/kg) may increase extracellular glutamate levels in the prefrontal cortex (Lorrain et al., 2003; Moghaddam et al., 1997), which may shift the balance between AMPA-to-NMDA receptors-mediated neurotransmission and contribute to the antidepressant-like effect of ketamine (Maeng et al., 2008; Zanos and Gould, 2018). Additionally, there is some evidence that the activation of AMPA receptors could increase anxiety (Alt et al., 2006; Andreasen et al., 2015; Fitzpatrick et al., 2016). Under these circumstances, it is possible that the activation of AMPA receptors mediated by increased ketamine induced-glutamate neurotransmission would increase anxiety, an effect that would counteract the possible anxiolytic-like effect of ketamine mediated by blocking the NMDA receptors. On the other hand, MK-801 increased extracellular glutamate levels in the prefrontal cortex only at high doses (0.3-0.6 mg/kg) (Roenker et al., 2012; Zuo et al., 2006). Consequently, a very low dose of MK-801 like the one used in our study (0.05 mg/kg) probably did not increase glutamate levels, which resulted in an anxiolytic-like effect mediated by the exclusive NMDA receptors blockade. This hypothesis, however, warrants further investigation and it is beyond the scope of the present study.

Meanwhile, a limitation of our study is that we evaluated ketamine and MK-801 effects only in one animal model related to PD. Given that, future studies should evaluate ketamine and MK-801 effects in other animal models of panic, such as the electrical stimulation of the dorsal periaqueductal grey matter (Moreira et al., 2013; Schenberg et al., 2001).

In summary, the present study provided an overview on the current state of knowledge about ketamine effects on anxiety/fear-related behaviors. At the same time, sub-anesthesic and anesthesic doses of ketamine had no effect on anxiety or panic-related behaviors in the ETM. The individual effects of R-ketamine and S-ketamine on anxiety/fear-related behaviors, including sex differences, should be further addressed.

## Acknowledgements

GPS received a master fellowship from Fundação de Apoio à Pesquisa e Inovação do Espírito Santo (FAPES) and received a doctoral research fellowship from Conselho Nacional de Desenvolvimento Científico e Tecnológico (CNPq) and now receives from Fundação de Amparo à Pesquisa do Estado de São Paulo (FAPESP; Grant#2018/12119-6; #2017/26815-1). SFSO and MMS received undergraduate research fellowships from UFES, DER is recipient of a fellowship from the Incentive Program to Post-Doctorate Attraction of the University of São Paulo, Brazil, and RA and SRLJ receive productivity fellowships from CNPq. This research did not receive any specific grant from funding agencies in the public, commercial, or not-for-profit sectors.

